# Small-molecule perturbation profiling reveals a mechanistic link between STING signaling and lipid metabolism in macrophages and dendritic cells

**DOI:** 10.1101/2025.04.25.650730

**Authors:** Lilah Gmyrek, Jianan Zhang, Maria Andrade, Kathryn R. Hillette, Xu Wu, Jin Mo Park

## Abstract

The DNA sensor cGAS and the signaling adaptor STING play a key role in the innate immune response to microbial and endogenous DNA in the cytoplasm. The cGAS-STING signaling pathway has evolved to promote immune defense and organismal fitness, yet its dysregulation can lead to chronic inflammation, autoimmunity, and neurodegeneration. Upon sensing double-stranded DNA, cGAS produces a cyclic dinucleotide second messenger that binds to STING in the endoplasmic reticulum. Ligand-bound STING translocates to the Golgi and activates the protein kinase TBK1 and the transcription factor IRF3, resulting in IRF3-driven interferon (IFN) gene transcription. This cascade of molecular events is mechanistically linked to intracellular lipid membrane dynamics and protein lipidation. To explore whether STING signaling is controlled by the availability and metabolic flux of cellular lipids, we screened small-molecule compounds targeting lipid metabolic pathways for their influence on STING agonist-responsive IFN induction. These screens identified inhibitors of the fatty acid synthase FAS and lipases as potent suppressors of STING signaling. An inhibitor of the cholesterol-esterifying enzyme SOAT1 enhanced STING-dependent IFN induction in mouse cells while attenuating it in human cells. Our findings reveal a connection between STING signaling and lipid metabolism and opportunities for expanding the toolbox for treating clinical conditions that arise from aberrant STING activity.

## Introduction

Stimulator of interferon genes (STING) is an endoplasmic reticulum (ER)-resident signaling adaptor and plays a central role in the innate immune response to cytoplasmic DNA (1–5). STING mediates intracellular signaling triggered by the double-stranded DNA sensor cyclic GMP–AMP synthase (cGAS), which recognizes microbial and endogenous DNA in a sequence-independent manner (6) and produces 2’3’-cyclic GMP-AMP (7–9). Upon binding to 2’3’-cyclic GMP-AMP (8–10), STING undergoes a series of molecular events leading to its activation— conformational changes and higher-order oligomerization (11, 12), ER-to-Golgi translocation (13, 14), and palmitoylation (15, 16). The active form of STING recruits the protein kinase TBK1 and the transcription factor IRF3 to its C-terminal tail (CTT) and mediates CTT and IRF3 phosphorylation by TBK1 (17–19). Phosphorylated IRF3 enters the nucleus and drives interferon (IFN) gene transcription.

STING signaling, when activated and controlled in a context-appropriate manner, contributes to immune defense against microbial pathogens and cancer. The importance of this protective function is highlighted by STING-dependent host resistance to herpes simplex virus-1: mice with a genetic ablation of STING fail to produce type I IFN upon exposure to this double-stranded DNA virus and succumb to its infection (5). STING-deficient mice also exhibit a defect in spontaneous and radiation-induced antitumor immune responses (20, 21). STING agonists with improved pharmacological properties have been developed for therapeutic activation of STING signaling.

Unrestrained STING activation and IFN induction in response to self-DNA can cause immunopathology, as seen in a wide range of genetic disorders and chronic diseases associated with inflammation, autoimmunity, and neurodegeneration. STING-associated vasculopathy with onset in infancy (SAVI) is a case in point: this autoinflammatory disorder with skin and lung manifestations arises from gain-of-function mutations in the STING gene (22). Aberrant mitochondrial retention in erythroid cells drives STING-dependent type I IFN production by macrophages in systemic lupus erythematosus (23). Mutations in the TDP43 and C9ORF72 genes, which are associated with amyotrophic lateral sclerosis and frontotemporal dementia, promote neurodegeneration through excessive STING activation. These mutations activate STING signaling by causing cytoplasmic TDP43 accumulation and TDP43-mediated mitochondrial DNA leakage (24) or by preventing C9ORF72-dependent lysosomal degradation of STING (25). STING activation by mitochondrial DNA in senescent cells has been shown to be a main driver of aging-related inflammation, neurodegeneration, and tissue dysfunction in mice (26). In addition to the well-established molecular events occurring within the cGAS-STING pathway, a diverse array of cellular processes and physiological conditions are expected to exert critical influence on STING activation and the resulting immune responses. Little information is available, however, regarding the wider molecular network in which STING serves its functions in health and disease.

Here, we discover novel interactions between STING signaling and lipid metabolism in macrophages and dendritic cells (DCs). Our study identifies small-molecules inhibitors of distinct lipid metabolic pathways that potently suppress or enhance IFN induction in STING agonist-treated cells. We show that perturbations in lipid metabolism can exert divergent effects on STING signaling in human and mouse cells, a phenomenon likely attributable to the difference of human and mouse STING proteins in their intrinsic capacity for lipid binding. These findings provide new insights into how STING transduces signals in the setting of active metabolism in living cells and point to pharmacologically actionable molecular processes operating at the nexus of immunity and lipid metabolism.

## Results

### STING activation and IFN induction in mouse and human cells treated with non-nucleotide agonists

We postulated an interplay between STING signaling and lipid metabolism given that STING signaling involves molecular trafficking and partitioning between membrane compartments, reversible long-chain fatty acid modification, and other processes controlled by the availability and flux of cellular lipid components (Fig. 1A). To explore the possibility of such interplay, we sought to determine whether pharmacological perturbation of lipid metabolic pathways alters STING-driven IFN induction. We established an assay system for this investigation using DC2.4, a mouse cell line with lineage and functional features of DCs (27), and 5,6-dimethylxanthenone-4-acetic acid (DMXAA; Fig. 1B), a non-nucleotide heterocyclic compound with mouse-specific STING agonist activity (28–30). Treatment with DMXAA induced STING and TBK1 phosphorylation as well as a change in the gel mobility of STING protein in cultured DC2.4 cells (Fig. 1C). Consistent with observations from several previous studies (4, 31, 32), STING protein from DMXAA-treated cells migrated as higher molecular weight forms during gel electrophoresis under a reducing and denaturing condition. This shift in gel mobility is thought to represent covalent modifications, reduction-resistant disulfide bridges, or other persistent molecular attributes associated with agonist-induced activation. DMXAA treatment also resulted in a drastic increase in the expression of the gene encoding IFN-β (*Ifnb1*), the flagship member of type I IFN (Fig. 1D). The strength of this induction was orders of magnitude greater than those observed with the toll-like receptor (TLR) agonists used for comparison (Fig. 1D).

**Figure 1.**
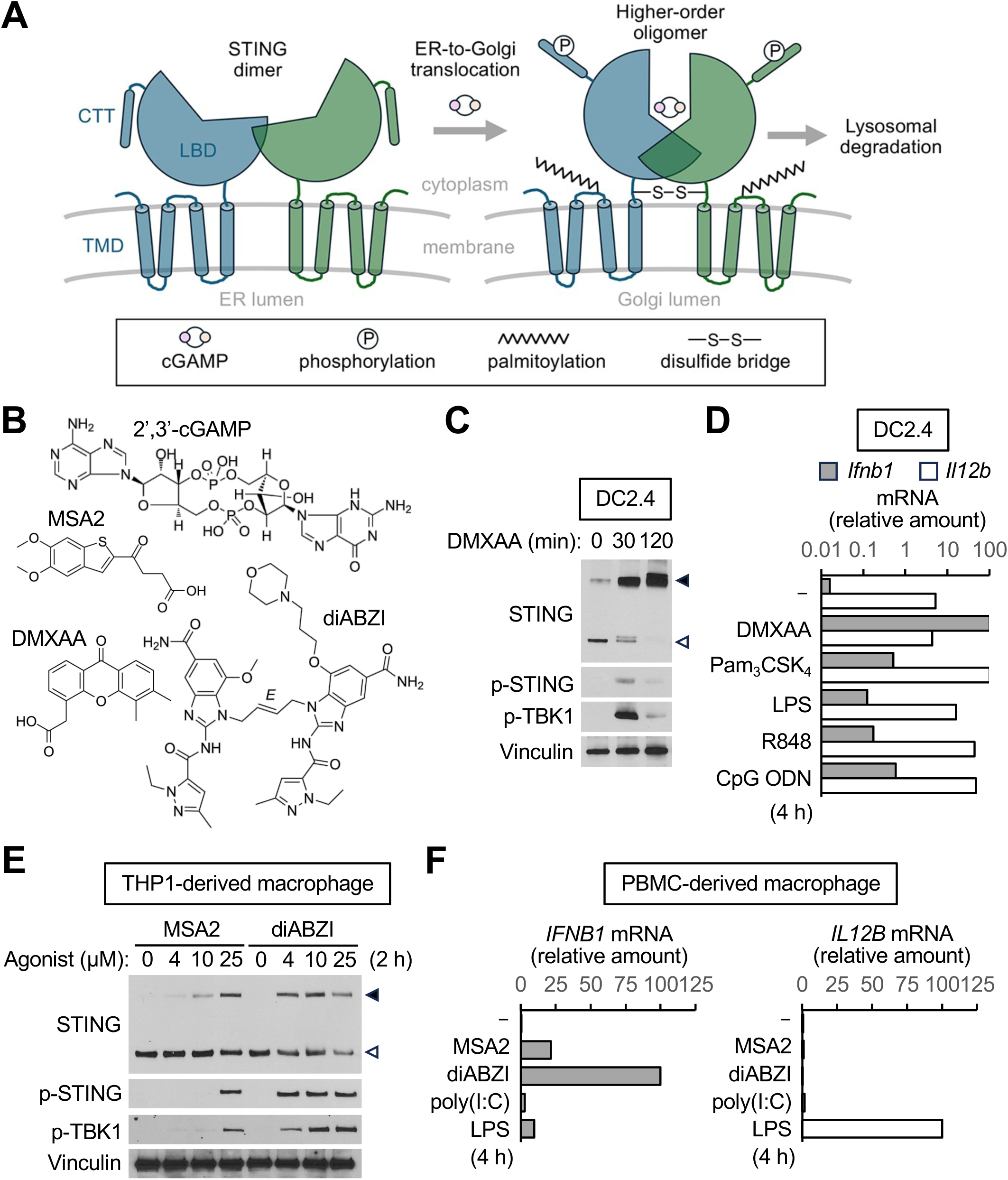
Assay systems established for perturbation of STING signaling in mouse and human cells. (**A**) Ligand-induced changes in STING protein structure and covalent modification. ER-resident STING exists as a dimer in the steady state. Upon ligand binding, STING forms oligomeric clusters in the Golgi compartment and undergoes other changes as indicated. Activated STING is degraded in lysosomes. TMD, transmembrane domain; LBD, ligand-binding domain; CTT, C-terminal tail. (**B**) STING agonists used in the study. (**C** and **D**) Immunoblot (**C**) and RT-qPCR (**D**) analysis of DC2.4 cells treated with DMXAA (50 μg/ml), Pam_3_CSK_4_ (1 μg/ml) lipopolysaccharide (LPS; 100 ng/ml), R848 (1 μM), or CpG oligodeoxynucleotide (ODN; 1 μM). (**E** and **F**) Immunoblot (**C**) and RT-qPCR (**D**) analysis of THP1-derived macrophages treated with MSA2 (25 μM or as indicated), diABZI (10 μM or as indicated), poly(I:C) (10 μg/ml), or LPS (100 ng/ml). p-, phosphorylated; filled arrowhead, high molecular weight forms of STING; open arrowhead, STING with an apparent size consistent with the calculated molecular weight.

Human and mouse STING differ in their expression pattern across cell types, affinity and responsiveness to cyclic dinucleotide (CDN) and non-nucleotide ligands, and ability to trigger downstream signaling events. We therefore set up another assay system for perturbation of human STING signaling using macrophages differentiated from the human monocytic cell line THP1 (33) and the cell-permeable non-nucleotide STING agonists MSA2 and diABZI (Fig. 1B), which were developed as cancer therapeutics (34, 35). Both agonists triggered STING signaling (Fig. 1E) and *IFNB1* induction (Fig. 1F) in macrophages derived from THP1 cells or human peripheral blood mononuclear cells (PBMCs). MSA2 and diABZI induced *IFNB1* expression with substantially higher intensities than the TLR agonists also examined (Fig. 1F). Together, these results showed that the two assay systems we established using mouse and human cells sensitively responded to STING agonists and could be used to assess the ability of lipid metabolic pathway inhibitors to alter STING signaling and STING-driven gene expression.

### Lipid metabolic pathway inhibitors suppressing or enhancing STING signaling

We examined a panel of small-molecule compounds blocking lipid metabolic pathways for their effects on DMXAA-responsive *Ifnb1* induction in DC2.4 cells. This panel of compounds included inhibitors of specific steps of lipid metabolism and covered its diverse aspects, ranging from fatty acid synthesis and oxidation to neutral lipid formation and breakdown. Four compounds were identified from this screen (Fig. 2A). Three of them—the lipase inhibitor CAY10499, the fatty acid synthase (FAS) inhibitor C75, and the glycolysis inhibitor 2- deoxyglucose—suppressed *Ifnb1* expression in DMXAA-treated cells. By contrast, the sterol O-acyltransferase-1 (SOAT1) inhibitor avasimibe enhanced *Ifnb1* induction in the same experimental condition. These inhibitors target enzymes involved in distinct anabolic and catabolic processes, yet they are expected to exert the shared effect of prompting the redistribution of neutral lipids, cholesterol in particular, between subcellular compartments.

**Figure 2.**
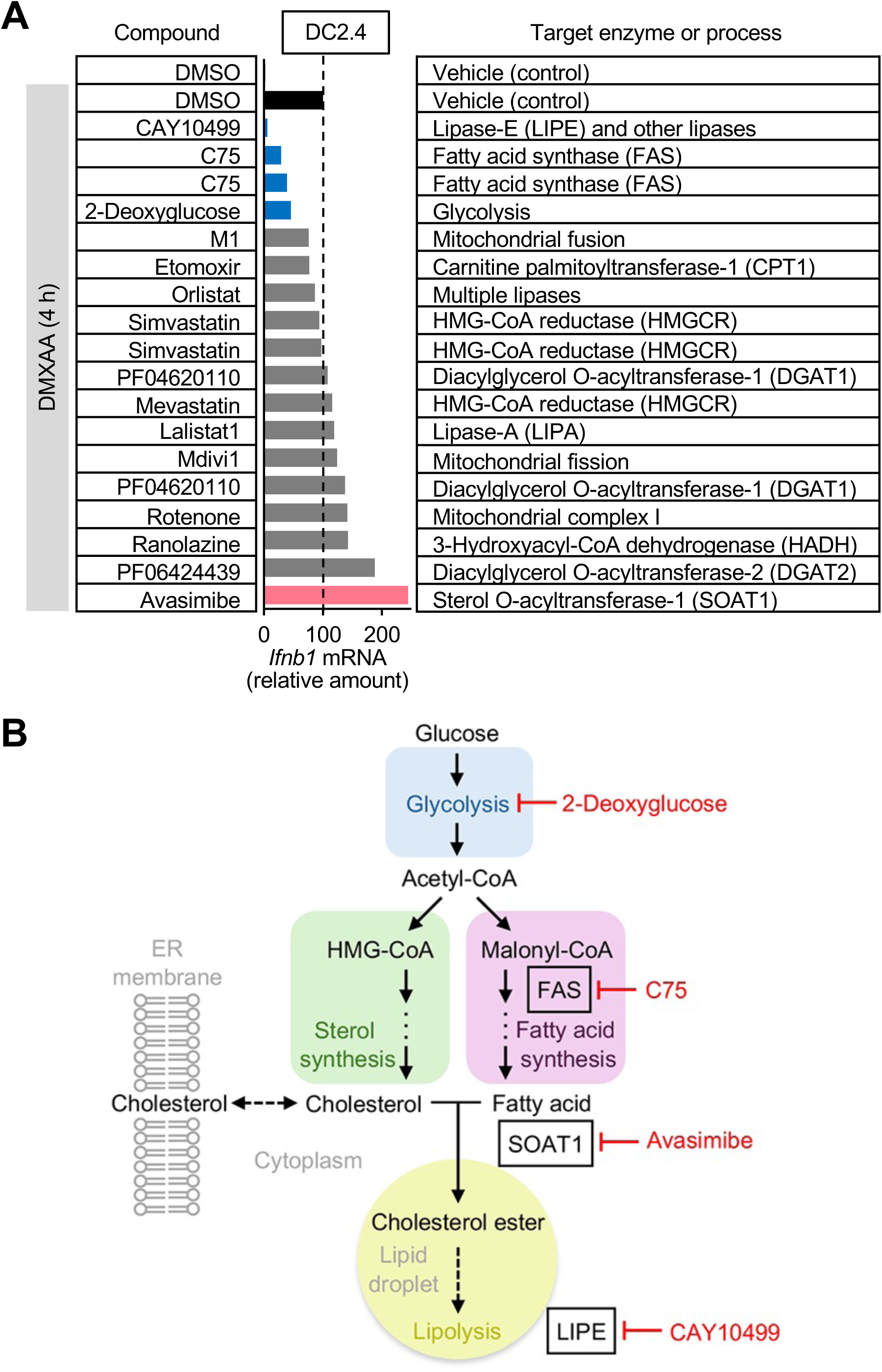
Small-molecule perturbation of STING agonist-responsive *Ifnb1* induction in mouse cells. (**A**) RT-qPCR analysis of DC2.4 cells treated with DMXAA (50 μg/ml) after preincubation with small-molecule compounds (concentrations indicated in Materials and Methods) for 30 min. (**B**) Small-molecule compounds suppressing or enhancing STING-driven *Ifnb1* induction and their target enzymes in lipid metabolic pathways.

SOAT1 is a key mediator of cholesterol repartitioning between the ER and cytoplasmic lipid droplets (Fig. 2B).

We performed an independent pharmacological perturbation profiling using THP1- derived macrophages; we examined the effects of lipid metabolic pathway inhibitors on MSA2- induced STING and TBK1 phosphorylation. CAY10499 emerged again as a potent suppressor of STING signaling in this screen and validation experiments (Fig. 4, A and B). The carnitine palmitoyltransferase-1 inhibitor etomoxir also suppressed STING signaling in THP1-derived macrophages (Fig. 3), whereas it had only a moderate effect on DMXAA-responsive *Ifnb1* induction in DC2.4 cells (Fig. 2A). Unlike their effect on *Ifnb1* induction in DC2.4 cells, C75 and 2-deoxyglucose left MSA2-triggered signaling intact in the screen using THP1-derived macrophages (Fig. 3). Subsequent validation experiments, however, revealed that C75 diminished diABZI-triggered signaling in macrophages derived from THP1 cells or primary human CD14^+^ monocytes (Fig. 4, A and B). Avasimibe, which enhanced *Ifnb1* induction in DMXAA-treated DC2.4 cells (Fig. 2A), attenuated MSA2- and diABZI-responsive phosphorylation events in THP1- and primary monocyte-derived macrophages in both discovery and validation experiments (Fig. 3; Fig. 4, A and B), a result indicative of its divergent effects on mouse and human STING signaling. The data obtained from the two independent screens led us to focus on C75, avasimibe, and CAY10499 for further investigation.

**Figure 3.**
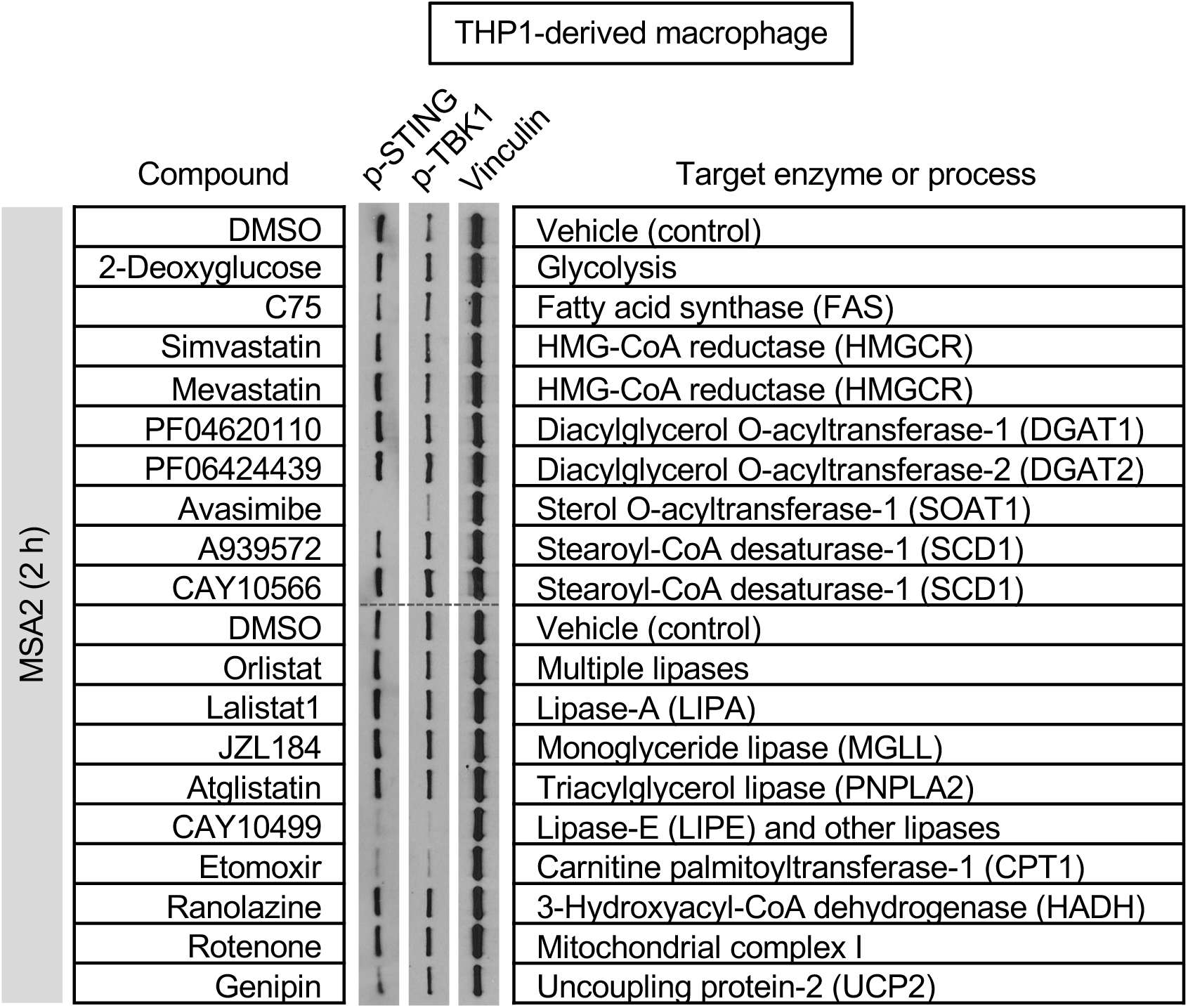
Small-molecule perturbation of STING agonist-responsive signaling in human cells. Immunoblot analysis of THP1-derived macrophages treated with MSA2 (25 μM) after preincubation with small-molecule compounds (concentrations indicated in Materials and Methods) for 30 min. p-, phosphorylated.

**Figure 4.**
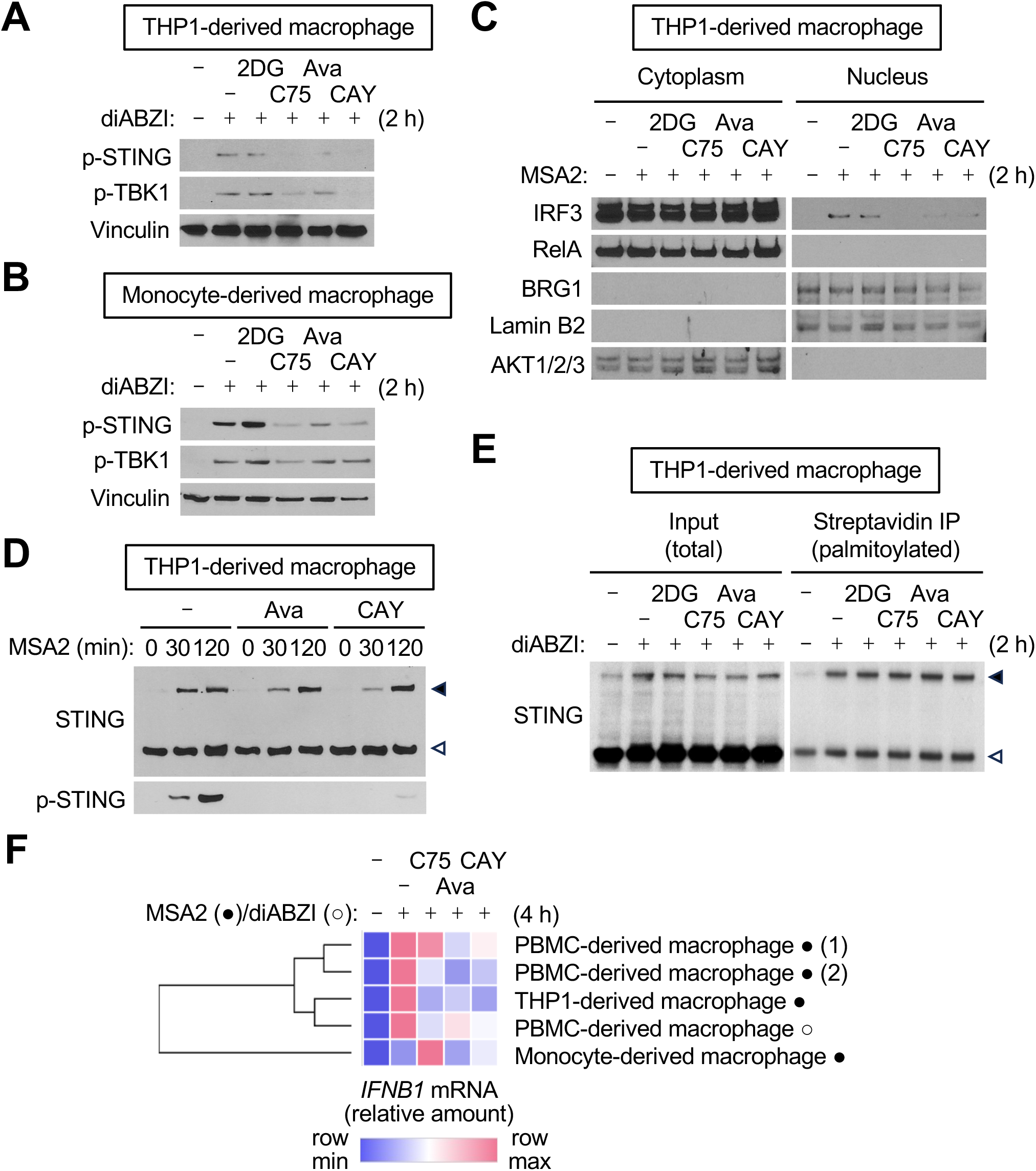
Effects of lipid metabolic pathway inhibitors on STING-driven signaling and *IFNB1* induction in human cells. (**A**-**E**) Immunoblot analysis of the indicated macrophage preparations treated with diABZI (10 μM) or MSA2 (25 μM) after preincubation with small-molecule compounds (concentrations indicated in Materials and Methods) for 30 min. Whole-cell lysates (**A**, **B**, **D**), cytoplasmic and nuclear fractions (**C**), and total and palmitoylated protein pools from whole-cell lysates (**E**) were used for analysis. (**F**) RT-qPCR analysis of the indicated macrophage preparations treated with diABZI (10 μM) or MSA2 (25 μM) after preincubation with small-molecule compounds for 30 min. Relative *IFNB1* mRNA amounts are presented on a color-coded scale shown at the bottom. p-, phosphorylated; 2DG, 2-deoxyglucose; Ava, avasimibe; CAY, CAY10499.

The entry of IRF3 into the nucleus occurred in THP1-derived macrophages after MSA2 treatment; C75, avasimibe, and CAY10499 blocked this translocation (Fig. 4C), showing that they exerted control across the full range of the STING-TBK1-IRF3 axis. Notably, high molecular weight forms of STING protein with altered gel mobility emerged in MSA2-treated cells even when its phosphorylation was prevented by avasimibe or CAY10499 (Fig. 4D). To determine whether the lipid metabolic pathway inhibitors affected STING palmitoylation, we labeled proteins in live THP1-derived macrophages with alkyne palmitic acid and isolated palmitoylated protein pools from their extracts by click chemistry-based biotin conjugation and streptavidin capture. Immunoblot analysis of the captured proteins revealed that none of the lipid metabolic pathway inhibitors examined reduced the extent of STING palmitoylation (Fig. 4E). Taken together, these findings suggest that STING adopts high molecular weight forms and becomes palmitoylated prior to or independently of its phosphorylation. C75, avasimibe, and CAY10499 likely interfered with signaling events upstream of TBK1-mediated phosphorylation of STING while sparing its palmitoylation and the molecular changes responsible for its gel mobility shift. The three small-molecule inhibitors suppressed STING agonist-responsive *IFNB1* induction in various human macrophage preparations (Fig. 4F).

### Conserved and species-specific effects of lipid metabolic pathway inhibitors on STING signaling

SOAT1 inhibition by avasimibe augmented STING agonist-responsive IFN induction in mouse cells (Fig. 2A) but attenuated it in human cells (Fig. 4F). Similarly, avasimibe enhanced STING agonist-triggered signaling events in mouse RAW264.7 macrophages (Fig. 5A) while dampening them in macrophages of human origin (Fig. 3; Fig. 4, A-C). The phosphorylation of TBK1 and IRF3 in RAW264.7 cells peaked at 30 min after DMXAA exposure and declined afterwards, but their phosphorylation was prolonged to at least 120 min with avasimibe treatment (Fig. 5A). In a similar setting, lipase inhibition by CAY10499 suppressed DMXAA-induced phosphorylation events, whereas lalistat1, an inhibitor specific to the lysosomal acid lipase LIPA, had no such effect (Fig. 5A). Avasimibe enhanced *Ifnb1* induction in various mouse macrophage and DC preparations (Fig. 5B). C75 and CAY10499, on the other hand, suppressed this STING-driven gene induction in mouse cells (Fig. 5B) as they did in human cells (Fig. 4F).

**Figure 5.**
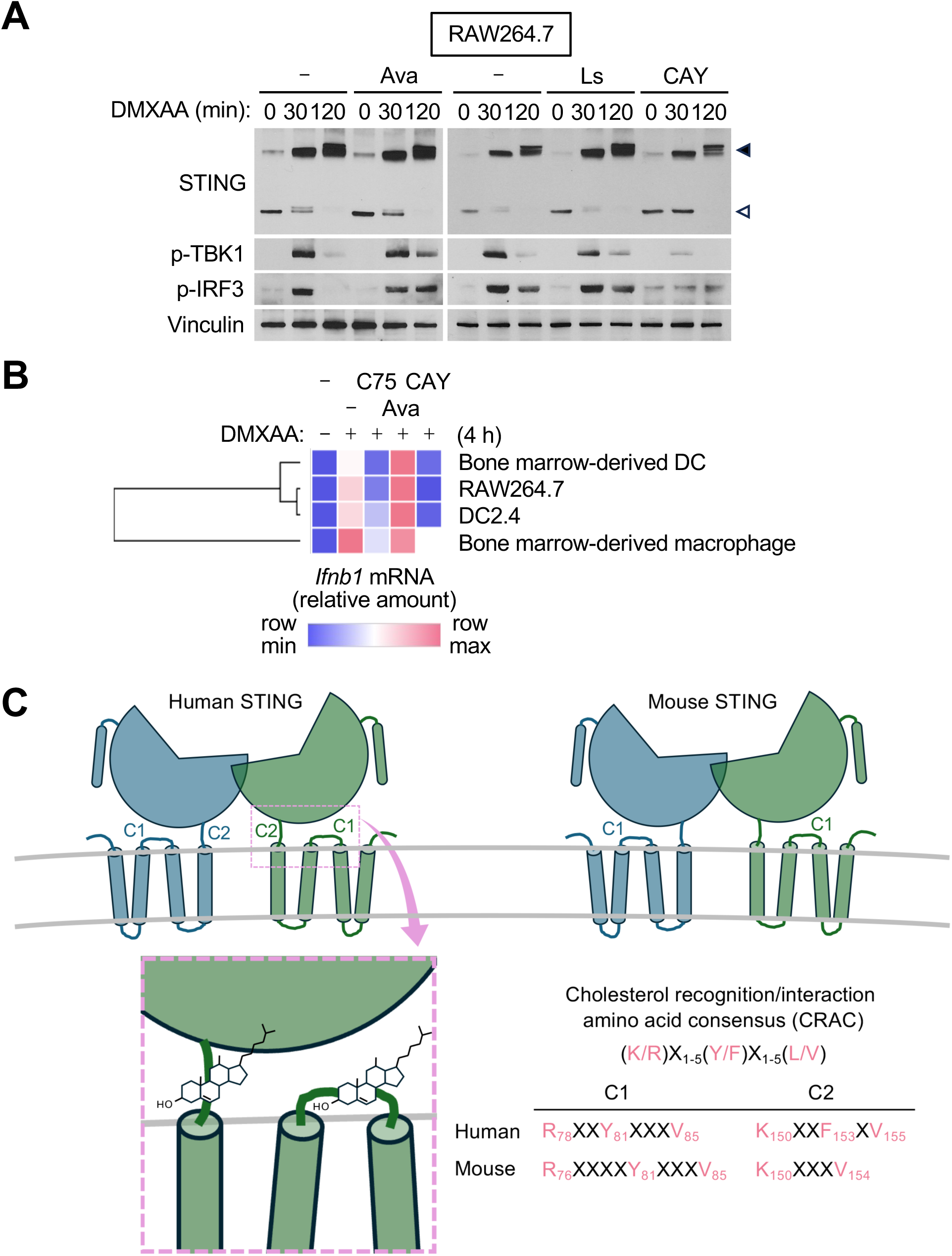
Effects of lipid metabolic pathway inhibitors on STING-driven signaling and *Ifnb1* induction in mouse cells. (**A**) Immunoblot analysis of RAW264.7 cells treated with DMXAA (50 μg/ml) after preincubation with small-molecule compounds (concentrations indicated in Materials and Methods) for 30 min. (**B**) RT-qPCR analysis of the indicated macrophage and DC preparations treated with DMXAA (50 μg/ml) after preincubation with small-molecule compounds for 30 min. Relative *Ifnb1* mRNA amounts are presented on a color-coded scale shown at the bottom. (**C**) CRAC sites, C1 and C2, in human and mouse STING proteins. CRAC sequence residues are indicated by the one-letter amino acid code, with X denoting any amino acid. p-, phosphorylated; Ava, avasimibe; Ls, lalistat1; CAY, CAY10499.

Human STING possesses two cholesterol recognition/interaction amino acid consensus (CRAC) sites, C1 and C2, within or near the transmembrane domain, which serve as functional cholesterol-interacting motifs (36). We analyzed the mouse STING polypeptide sequence and found that it retained a sequence motif corresponding to C1 but did not have a C2 equivalent that conforms to the CRAC sequence (Fig. 5C). Non-identical CRAC site configurations in human and mouse STING are thought to make the two proteins interact with cholesterol differently and underlie the contrasting effects of avasimibe on STING signaling and IFN induction in the two species.

## Discussion

Our findings link STING-dependent innate immune responses to lipid metabolism. The results of our pharmacological perturbation profiling and validation experiments suggest that fatty acid synthesis, cholesterol esterification, and neutral lipid breakdown—mediated by FAS, SOAT1, and lipases, respectively—promote or regulate STING signaling. Inhibitors of these enzymes, C75, avasimibe, and CAY10499, exert their effects on STING signaling in macrophages and DCs. These inhibitors are thought to alter the availability and metabolic flux of fatty acids and neutral lipids, thereby reshaping the molecular milieu in which STING activation and inactivation occur. Both FAS and lipases are involved in metabolic processes that generate free fatty acids. Their inhibition by C75 and CAY10499, however, did not affect STING palmitoylation in our analysis. It remains to be determined whether fatty acid availability is a critical factor limiting the palmitoylation of other proteins or controlling other signaling events in the STING pathway. Lipase-E (LIPE; also known as hormone-sensitive lipase) is one of the enzymes responsible for the breakdown of neutral lipids sequestered in cytoplasmic lipid droplets. LIPE is sensitive to inhibition by CAY10499 but not by lalistat1. Lipid droplets in macrophages have been implicated in innate immune function and antimicrobial defense (37). Therefore, LIPE may promote STING signaling as a mediator of lipid droplet turnover.

SOAT1 is an ER-resident enzyme required for the esterification of free cholesterol, hence its sequestration in the form of cholesterol ester in cytoplasmic lipid droplets. SOAT1 inhibition reverses this process, resulting in a greater availability of cholesterol for other subcellular compartments including the ER. Cholesterol in the ER membrane binds to STING and prevents its trafficking and activation (36). Human STING is amenable to control by ER cholesterol, as it has two functional CRAC sites. SOAT1 inhibitor-induced cholesterol repartitioning and the resulting overabundance of ER cholesterol create a condition that suppresses the activation of human STING. Mouse STING is presumably insensitive to this control, as it lacks a fully conserved cholesterol-interacting motif configuration. In contrast to cholesterol in the ER, lysosomal cholesterol enhances STING activation by disrupting a process crucial for restraining STING signaling: lysosomal degradation of activated STING (38). Cholesterol in lysosomes suppresses vacuolar ATPase-dependent luminal acidification, a prerequisite for not merely STING proteolysis but also the hydrolytic activity of these organelles in general (39, 40). Lysosomal cholesterol exerts this action independently of functional CRAC sites in the proteins targeted for degradation. We postulate that SOAT1 inhibition alters lysosomal cholesterol content and prevents timely termination of STING signaling. In line with this idea, we observed that SOAT1 inhibition by avasimibe prolonged TBK1 and IRF3 activation and augmented *Ifnb1* induction in STING agonist-treated mouse cells. The divergent effects of SOAT1 inhibition on human and mouse STING signaling may be the consequence of the difference in STING- cholesterol interactions in the two species as well as the opposite actions of ER versus lysosomal cholesterol on STING activation.

This study introduces new approaches to manipulating STING signaling for clinical benefit. Inhibitors of FAS, SOAT1, and lipases have been studied in clinical trials for indications including cancer, atherosclerosis, and dyslipidemia. Repurposed and newly developed drugs targeting these enzymes may be effective at suppressing aberrant STING activity in human diseases and potentiating STING-driven antitumor immunity. Furthermore, diet and lifestyle changes that lead to reprogramming in lipid metabolism have the potential to modulate STING signaling and have an impact on diseases and therapy responses where STING plays an important role.

## Materials and Methods

### Primary cells and cell lines

DC2.4 and RAW264.7 cells were cultured in Gibco DMEM medium (Thermo Fisher Scientific) supplemented with FBS (10%), penicillin (100 U/ml), and streptomycin (100 μg/ml). Mouse C57BL/6 bone marrow cells were cultured in Gibco DMEM medium (Thermo Fisher Scientific) supplemented with FBS, sodium pyruvate (1 mM), L-glutamine (2 mM), penicillin, and streptomycin. Mouse bone marrow cells were treated with mouse macrophage-colony stimulating factor (10 ng/ml, PeproTech) for 7 days and with mouse FMS-like tyrosine kinase-3 ligand (10 ng/ml, PeproTech) for 9 days to induce their differentiation into macrophages and DCs, respectively. THP-1 cells were cultured in Gibco RPMI 1640 medium (Thermo Fisher Scientific) supplemented with FBS, penicillin, and streptomycin. THP1 cells were treated with phorbol 12-myristate 13-acetate (100 nM; Sigma-Aldrich) for 3 days to induce their differentiation into macrophages. Human PBMCs and CD14^+^ monocytes were cultured in RPMI 1640 medium (Life Technologies) supplemented with FBS, sodium pyruvate, L-glutamine, β-mercaptoethanol (50 uM), penicillin, and streptomycin. Human PBMCs and CD14^+^ monocytes were treated with human macrophage-colony stimulating factor (10 ng/ml, PeproTech) for 7 days to induce their differentiation into macrophages.

### Immunostimulatory agents and small-molecule compounds

Cells were treated with the following immunostimulatory agents to activate STING or TLR signaling: DMXAA, Pam_3_CSK_4_, poly(I:C), R848, and CpG ODN (ODN1668; all from InvivoGen); MSA2 and diABZI (both from Selleck Chemicals); and LPS (Sigma-Aldrich). Cells were treated with the following small-molecule compounds for perturbation of lipid metabolic pathways: C75, PF04620110, PF06424439, avasimibe, etomoxir, ranolazine, olistat, JZL184, atglistatin, mevastatin, simvastatin, genipin, Mdivi1, and A939572 (all from Selleck Chemical); 2-deoxyglucose, rotenone, and M1 (all from Sigma-Aldrich); and lalistat1, CAY10499, and CAY10566 (all from Cayman Chemical). All compounds were added to medium at a final concentration of 20 μM except for 2-deoxyglucose (1.5 mM), etomoxir (200 μM), ranolazine (200 μM), mevastatin (50 μM), simvastatin (10 μM), and rotenone (1 μM).

### RNA isolation, cDNA synthesis, and RT-qPCR

For RT-qPCR analysis, total RNA isolated using the Trizol Reagent (Thermo Fisher Scientific) were subjected to cDNA synthesis using the SuperScript IV VILO Master Mix (Thermo Fisher Scientific) and PCR using the SYBR Green PCR Master Mix (Thermo Fisher Scientific) and the following gene-specific primers: *Ppia*, 5’- atggtcaaccccaccgtgt-3’ (forward) and 5’-ttcttgctgtctttggaactttgtc-3’ (reverse); *Ifnb1*, 5’- cgtgggagatgtcctcaact-3’ (forward) and 5’-acctttgcaccctccagtaa-3’ (reverse); *Il12b*, 5’- atccagcgcaagaaagaaaa-3’ (forward) and 5’-ggaacgcacctttctggtta-3’ (reverse); *GAPDH*, 5’- cgtggaaggactcatgacca-3’ (forward) and 5’-gccatcacgccacagtttc-3’ (reverse); *IFNB1*, 5’- tgggaggattctgcattacc-3’ (forward) and 5’-ctatggtccaggcacagtga-3’ (reverse); and *IL12B*, 5’- aacttgcagctgaagccatt-3’ (forward) and 5’-tactcccagctgacctccac-3’ (reverse). Heat maps and dendrograms visualizing gene transcript abundances and hierarchical mRNA profile clustering, respectively, were generated using Morpheus software (Broad Institute). Hierarchical clustering was performed using the average linkage method and one minus Pearson’s correlation as the distance metric.

### Protein isolation and immunoblotting

To prepare whole-cell lysates, cells were incubated in cell lysis buffer (0.2 ml for 10^6^ cells; Cell Signaling Technology) containing phenylmethylsulfonyl fluoride (PMSF; 1 mM) for 5 min at 4°C. Lysed cells in the buffer were centrifuged 5 min at 16,000 g. The supernatants were collected for use as whole-cell lysates. To prepare cytoplasmic and nuclear fractions, cells were incubated in buffer L1 (1 ml for 10^7^ cells; 50 mM Tris-chloride [pH 8.0], 2 mM EDTA, 0.1% Nonidet P-40, 10% glycerol, 25 mM β- glycerophosphate, and 1 mM PMSF) for 5 min at 4°C. Intact nuclei and cell debris in the buffer were centrifuged 5 min at 4,500 g. The supernatants were stored for use as cytoplasmic fractions. The pelleted nuclei were rinsed with buffer L1 (1 ml for nuclei equivalent to 10^7^ cells) and resuspended in cell lysis buffer containing PMSF (0.2 ml for nuclei equivalent to 10^7^ cells), incubated for 5 min at 4°C, and centrifuged 5 min at 16,000 g. The supernatants were collected for use as nuclear fractions. Antibodies specific to the following proteins were used in immunoblotting after 1:1000 dilution: p-STING (human, 19781), p-STING (mouse, 72971), STING (13647), and p-TBK1 (5483; all from Cell Signaling Technology); AKT1/2/3 (sc-8312), BRG1 (sc-10768), RelA (sc-372), and vinculin (sc-73614; all from Santa Cruz Biotechnology); and IRF3 (51-3200) and lamin B2 (33-2100; both from Thermo Fisher Scientific).

### Protein palmitoylation analysis

Cells were incubated with alkyne palmitic acid (50 μM) in medium for 16 h before treatment with lipid metabolic pathway inhibitors and MSA2. The cells were incubated in RIPA buffer (0.1 ml for 10^6^ cells; 50 mM Tris-chloride [pH 7.4], 150 mM sodium chloride, 0.2% sodium dodecyl sulfate, 1% Triton X-100) for 30 min at 4°C and sonicated. Lysed cells in the buffer were centrifuged 5 min at 16,000 g. The supernatants were subjected to a click reaction (100 μM biotin azide, 200 μM TBTA, 1 mM copper[II] sulfate, 1 mM TCEP) for 1 h at room temperature. Proteins in the reactions were precipitated, resuspended in 1% SDS buffer (50 mM TEA [pH 7.4], 150 mM sodium chloride, 1% sodium dodecyl sulfate), and diluted with 1% Triton X-100 buffer (50 mM TEA [pH 7.4], 150 mM sodium chloride, 1% Triton X-100) to reduce sodium dodecyl sulfate concentration to 0.2%. Alkyne-palmitoylated proteins in the resuspended fractions were captured by incubation with streptavidin beads for 3 h at room temperature, eluted from the beads using the SDS gel loading buffer, and analyzed by immunoblotting.

## Acknowledgements

This study was supported by the US Department of Defense grant W81XWH-21-1-0074 (J.M.P.).

## Abbreviations

CDN: cyclic dinucleotide
cGAS: cyclic GMP-AMP synthase
CRAC: cholesterol recognition/interaction amino acid consensus
CTT: C-terminal tail
DC: dendritic cell
DMXAA: 5,6-dimethylxanthenone-4-acetic acid
ER: endoplasmic reticulum
FAS: fatty acid synthase
IFN: interferon
IRF3: interferon regulatory factor 3
LIPE: lipase-E
LPS: lipopolysaccharide
ODN: oligodeoxynucleotide
RT-qPCR: real-time quantitative polymerase chain reaction
SOAT1: sterol O-acyltransferase-1
STING: stimulator of interferon genes
TBK1: tank-binding kinase 1
TLR: toll-like receptor

## References

1. Jin L, Waterman PM, Jonscher KR, Short CM, Reisdorph NA, Cambier JC. MPYS, a novel membrane tetraspanner, is associated with major histocompatibility complex class II and mediates transduction of apoptotic signals. Mol Cell Biol. 2008;28(16):5014–5026.

2. Ishikawa H, Barber GN. STING is an endoplasmic reticulum adaptor that facilitates innate immune signalling. Nature. 2008;455(7213):674-678.

3. Zhong B, Yang Y, Li S, Wang YY, Li Y, Diao F, Lei C, He X, Zhang L, Tien P, Shu HB. The adaptor protein MITA links virus-sensing receptors to IRF3 transcription factor activation. Immunity. 2008;29(4):538–550.

4. Sun W, Li Y, Chen L, Chen H, You F, Zhou X, Zhou Y, Zhai Z, Chen D, Jiang Z. ERIS, an endoplasmic reticulum IFN stimulator, activates innate immune signaling through dimerization. Proc Natl Acad Sci U S A. 2009;106(21):8653–8658.

5. Ishikawa H, Ma Z, Barber GN. STING regulates intracellular DNA-mediated, type I interferon-dependent innate immunity. Nature. 2009;461(7265):788-792.

6. Civril F, Deimling T, de Oliveira Mann CC, Ablasser A, Moldt M, Witte G, Hornung V, Hopfner KP. Structural mechanism of cytosolic DNA sensing by cGAS. Nature. 2013;498(7454):332-337.

7. Sun L, Wu J, Du F, Chen X, Chen ZJ. Cyclic GMP-AMP synthase is a cytosolic DNA sensor that activates the type I interferon pathway. Science. 2013;339(6121):786-791.

8. Ablasser A, Goldeck M, Cavlar T, Deimling T, Witte G, Röhl I, Hopfner KP, Ludwig J, Hornung V. cGAS produces a 2’-5’-linked cyclic dinucleotide second messenger that activates STING. Nature. 2013;498(7454):380-384.

9. Diner EJ, Burdette DL, Wilson SC, Monroe KM, Kellenberger CA, Hyodo M, Hayakawa Y, Hammond MC, Vance RE. The innate immune DNA sensor cGAS produces a noncanonical cyclic dinucleotide that activates human STING. Cell Rep. 2013;3(5):1355–1361.

10. Wu J, Sun L, Chen X, Du F, Shi H, Chen C, Chen ZJ. Cyclic GMP-AMP is an endogenous second messenger in innate immune signaling by cytosolic DNA. Science. 2013;339(6121):826-830.

11. Shang G, Zhang C, Chen ZJ, Bai XC, Zhang X. Cryo-EM structures of STING reveal its mechanism of activation by cyclic GMP-AMP. Nature. 2019;567(7748):389-393.

12. Ergun SL, Fernandez D, Weiss TM, Li L. STING Polymer Structure Reveals Mechanisms for Activation, Hyperactivation, and Inhibition. Cell. 2019;178(2):290–301.e10.

13. Gui X, Yang H, Li T, Tan X, Shi P, Li M, Du F, Chen ZJ. Autophagy induction via STING trafficking is a primordial function of the cGAS pathway. Nature. 2019;567(7747):262-266.

14. Dobbs N, Burnaevskiy N, Chen D, Gonugunta VK, Alto NM, Yan N. STING Activation by Translocation from the ER Is Associated with Infection and Autoinflammatory Disease. Cell Host Microbe. 2015;18(2):157–168.

15. Mukai K, Konno H, Akiba T, Uemura T, Waguri S, Kobayashi T, Barber GN, Arai H, Taguchi T. Activation of STING requires palmitoylation at the Golgi. Nat Commun. 2016;7:11932.

16. Haag SM, Gulen MF, Reymond L, Gibelin A, Abrami L, Decout A, Heymann M, van der Goot FG, Turcatti G, Behrendt R, Ablasser A. Targeting STING with covalent small-molecule inhibitors. Nature. 2018;559(7713):269-273.

17. Liu S, Cai X, Wu J, Cong Q, Chen X, Li T, Du F, Ren J, Wu YT, Grishin NV, Chen ZJ. Phosphorylation of innate immune adaptor proteins MAVS, STING, and TRIF induces IRF3 activation. Science. 2015;347(6227):aaa2630.

18. Zhang C, Shang G, Gui X, Zhang X, Bai XC, Chen ZJ. Structural basis of STING binding with and phosphorylation by TBK1. Nature. 2019;567(7748):394-398.

19. Yum S, Li M, Fang Y, Chen ZJ. TBK1 recruitment to STING activates both IRF3 and NF- κB that mediate immune defense against tumors and viral infections. Proc Natl Acad Sci U S A. 2021;118(14):e2100225118.

20. Woo SR, Fuertes MB, Corrales L, Spranger S, Furdyna MJ, Leung MY, Duggan R, Wang Y, Barber GN, Fitzgerald KA, Alegre ML, Gajewski TF. STING-dependent cytosolic DNA sensing mediates innate immune recognition of immunogenic tumors. Immunity. 2014;41(5):830–842.

21. Deng L, Liang H, Xu M, Yang X, Burnette B, Arina A, Li XD, Mauceri H, Beckett M, Darga T, Huang X, Gajewski TF, Chen ZJ, Fu YX, Weichselbaum RR. STING-Dependent Cytosolic DNA Sensing Promotes Radiation-Induced Type I Interferon-Dependent Antitumor Immunity in Immunogenic Tumors. Immunity. 2014;41(5):843–852.

22. Liu Y, Jesus AA, Marrero B, Yang D, Ramsey SE, Sanchez GAM, Tenbrock K, Wittkowski H, Jones OY, Kuehn HS, Lee CR, DiMattia MA, Cowen EW, Gonzalez B, Palmer I, DiGiovanna JJ, Biancotto A, Kim H, Tsai WL, Trier AM, Huang Y, Stone DL, Hill S, Kim HJ, St Hilaire C, Gurprasad S, Plass N, Chapelle D, Horkayne-Szakaly I, Foell D, Barysenka A, Candotti F, Holland SM, Hughes JD, Mehmet H, Issekutz AC, Raffeld M, McElwee J, Fontana JR, Minniti CP, Moir S, Kastner DL, Gadina M, Steven AC, Wingfield PT, Brooks SR, Rosenzweig SD, Fleisher TA, Deng Z, Boehm M, Paller AS, Goldbach-Mansky R. Activated STING in a vascular and pulmonary syndrome. N Engl J Med. 2014;371(6):507–518.

23. Caielli S, Cardenas J, de Jesus AA, Baisch J, Walters L, Blanck JP, Balasubramanian P, Stagnar C, Ohouo M, Hong S, Nassi L, Stewart K, Fuller J, Gu J, Banchereau JF, Wright T, Goldbach-Mansky R, Pascual V. Erythroid mitochondrial retention triggers myeloid-dependent type I interferon in human SLE. Cell. 2021;184(17):4464–4479.e19.

24. Yu CH, Davidson S, Harapas CR, Hilton JB, Mlodzianoski MJ, Laohamonthonkul P, Louis C, Low RRJ, Moecking J, De Nardo D, Balka KR, Calleja DJ, Moghaddas F, Ni E, McLean CA, Samson AL, Tyebji S, Tonkin CJ, Bye CR, Turner BJ, Pepin G, Gantier MP, Rogers KL, McArthur K, Crouch PJ, Masters SL. TDP-43 Triggers Mitochondrial DNA Release via mPTP to Activate cGAS/STING in ALS. Cell. 2020;183(3):636–649.e18.

25. McCauley ME, O’Rourke JG, Yáñez A, Markman JL, Ho R, Wang X, Chen S, Lall D, Jin M, Muhammad AKMG, Bell S, Landeros J, Valencia V, Harms M, Arditi M, Jefferies C, Baloh RH. C9orf72 in myeloid cells suppresses STING-induced inflammation. Nature. 2020;585(7823):96-101.

26. Gulen MF, Samson N, Keller A, Schwabenland M, Liu C, Glück S, Thacker VV, Favre L, Mangeat B, Kroese LJ, Krimpenfort P, Prinz M, Ablasser A. cGAS-STING drives ageing-related inflammation and neurodegeneration. Nature. 2023;620(7973):374-380.

27. Shen Z, Reznikoff G, Dranoff G, Rock KL. Cloned dendritic cells can present exogenous antigens on both MHC class I and class II molecules. J Immunol. 1997;158(6):2723–2730.

28. Prantner D, Perkins DJ, Lai W, Williams MS, Sharma S, Fitzgerald KA, Vogel SN. 5,6- Dimethylxanthenone-4-acetic acid (DMXAA) activates stimulator of interferon gene (STING)-dependent innate immune pathways and is regulated by mitochondrial membrane potential. J Biol Chem. 2012;287(47):39776–39788.

29. Conlon J, Burdette DL, Sharma S, Bhat N, Thompson M, Jiang Z, Rathinam VA, Monks B, Jin T, Xiao TS, Vogel SN, Vance RE, Fitzgerald KA. Mouse, but not human STING, binds and signals in response to the vascular disrupting agent 5,6-dimethylxanthenone-4-acetic acid. J Immunol. 2013;190(10):5216–5225.

30. Kim S, Li L, Maliga Z, Yin Q, Wu H, Mitchison TJ. Anticancer flavonoids are mouse-selective STING agonists. ACS Chem Biol. 2013;8(7):1396–1401.

31. Tsuchida T, Zou J, Saitoh T, Kumar H, Abe T, Matsuura Y, Kawai T, Akira S. The ubiquitin ligase TRIM56 regulates innate immune responses to intracellular double-stranded DNA. Immunity. 2010;33(5):765–776.

32. Wang X, Majumdar T, Kessler P, Ozhegov E, Zhang Y, Chattopadhyay S, Barik S, Sen GC. STING Requires the Adaptor TRIF to Trigger Innate Immune Responses to Microbial Infection. Cell Host Microbe. 2016;20(3):329–341.

33. Tsuchiya S, Yamabe M, Yamaguchi Y, Kobayashi Y, Konno T, Tada K. Establishment and characterization of a human acute monocytic leukemia cell line (THP-1). Int J Cancer. 1980;26(2):171–176.

34. Pan BS, Perera SA, Piesvaux JA, Presland JP, Schroeder GK, Cumming JN, Trotter BW, Altman MD, Buevich AV, Cash B, Cemerski S, Chang W, Chen Y, Dandliker PJ, Feng G, Haidle A, Henderson T, Jewell J, Kariv I, Knemeyer I, Kopinja J, Lacey BM, Laskey J, Lesburg CA, Liang R, Long BJ, Lu M, Ma Y, Minnihan EC, O’Donnell G, Otte R, Price L, Rakhilina L, Sauvagnat B, Sharma S, Tyagarajan S, Woo H, Wyss DF, Xu S, Bennett DJ, Addona GH. An orally available non-nucleotide STING agonist with antitumor activity. Science. 2020;369(6506):eaba6098.

35. Ramanjulu JM, Pesiridis GS, Yang J, Concha N, Singhaus R, Zhang SY, Tran JL, Moore P, Lehmann S, Eberl HC, Muelbaier M, Schneck JL, Clemens J, Adam M, Mehlmann J, Romano J, Morales A, Kang J, Leister L, Graybill TL, Charnley AK, Ye G, Nevins N, Behnia K, Wolf AI, Kasparcova V, Nurse K, Wang L, Puhl AC, Li Y, Klein M, Hopson CB, Guss J, Bantscheff M, Bergamini G, Reilly MA, Lian Y, Duffy KJ, Adams J, Foley KP, Gough PJ, Marquis RW, Smothers J, Hoos A, Bertin J. Design of amidobenzimidazole STING receptor agonists with systemic activity. Nature. 2018;564(7736):439-443.

36. Zhang BC, Laursen MF, Hu L, Hazrati H, Narita R, Jensen LS, Hansen AS, Huang J, Zhang Y, Ding X, Muyesier M, Nilsson E, Banasik A, Zeiler C, Mogensen TH, Etzerodt A, Agger R, Johannsen M, Kofod-Olsen E, Paludan SR, Jakobsen MR. Cholesterol-binding motifs in STING that control endoplasmic reticulum retention mediate anti-tumoral activity of cholesterol-lowering compounds. Nat Commun. 2024;15(1):2760.

37. Bosch M, Sánchez-Álvarez M, Fajardo A, Kapetanovic R, Steiner B, Dutra F, Moreira L, López JA, Campo R, Marí M, Morales-Paytuví F, Tort O, Gubern A, Templin RM, Curson JEB, Martel N, Català C, Lozano F, Tebar F, Enrich C, Vázquez J, Del Pozo MA, Sweet MJ, Bozza PT, Gross SP, Parton RG, Pol A. Mammalian lipid droplets are innate immune hubs integrating cell metabolism and host defense. Science. 2020;370(6514):eaay8085.

38. Chu TT, Tu X, Yang K, Wu J, Repa JJ, Yan N. Tonic prime-boost of STING signalling mediates Niemann-Pick disease type C. Nature. 2021;596(7873):570-575.

39. Cox BE, Griffin EE, Ullery JC, Jerome WG. Effects of cellular cholesterol loading on macrophage foam cell lysosome acidification. J Lipid Res. 2007;48(5):1012–1021.

40. Jerome WG, Cox BE, Griffin EE, Ullery JC. Lysosomal cholesterol accumulation inhibits subsequent hydrolysis of lipoprotein cholesteryl ester. Microsc Microanal. 2008;14(2):138–149.

